# A new threat identified in the use of SDHIs pesticides targeting the mitochondrial succinate dehydrogenase enzyme

**DOI:** 10.1101/289058

**Authors:** Paule Bénit, Sylvie Bortoli, Laurence Huc, Manuel Schiff, Anne-Paule Gimenez-Roqueplo, Malgorzata Rak, Pierre Gressens, Judith Favier, Pierre Rustin

## Abstract

Succinate dehydrogenase inhibitors (SDHIs) are now widely used worldwide as fungicides to limit the proliferation of molds in cereal crops, or to better preserve fruits, vegetables, and seeds from these molds, as well as to facilitate the lawn care for public spaces and golf courses. According to the companies that produce them, the SDHIs quite specifically inhibit the activity of the succinate dehydrogenase in the molds. We here establish that these inhibitors readily inhibit the earthworm and the human enzyme, raising a new concern on the danger of their large scale utilization in agriculture. This is all the more worrying as we know that the loss of function, partial or total, of the SDH activity caused by genetic variants causes severe human neurological diseases, or leads to the development of tumors and/or cancers.

---

The succinate dehydrogenase (SDH), also known as respiratory chain complex II, is a universal and key component of the mitochondrial respiratory chain. As part of the Krebs cycle, this enzyme oxidizes succinate into fumarate with the concomitant reduction of respiratory chain ubiquinone (Fig 1). In human, germline mutations in one of the four genes encoding the SDH subunits (SDH A-D) result in either early onset severe neurological deterioration in childhood (*1*) or paragangliomas (tumors affecting the parasympathetic nervous system in the head and neck, or the extra-adrenal sympathetic nervous system in thoracic-abdominal or pelvic areas) and pheochromocytomas (tumors arising from the adrenal medulla) (*2*). A predisposition to the development of gastrointestinal stromal tumors (GIST), as well as to rare cases of kidney cancers, has also been observed (*3*). Noticeably, in human cultured cells, a partial defect of the SDH does not necessarily result in cell death (*4*), and tumor formation in affected individuals is not associated with random mutations as classically detected by genotoxicity tests. Tumor formation rather results from epigenetics modifications, which have been shown to be a long-term consequence of succinate accumulation, acting as an oncometabolite (*5*). We recently became aware of the large-scale utilization of inhibitors of the enzyme as fungicides, some of these acclaimed for their remanence. Nowadays, the so-called SDHIs (standing for SDH inhibitors) are recommended, alone or in association, to fight fungi proliferation in a number of situations, including the preservation of cereal crops, fruits, vegetables, and seeds, as well as for lawn care of public and golf lawns. The widespread use of SDHIs, particularly massive from 2013-2014 in France, came after the progressive withdrawal of several molecules targeting other components of the respiratory chain due to belatedly recognized hazard and/or appearance of fungi resistance to these drugs (*6*). Strikingly, we observed that the catalytic pocket of the succinate dehydrogenase targeted by the SDHIs is highly conserved between species, making doubtful an eventual specificity towards the fungi enzyme advocated by some of the companies producing these molecules.

**Figure 1.**
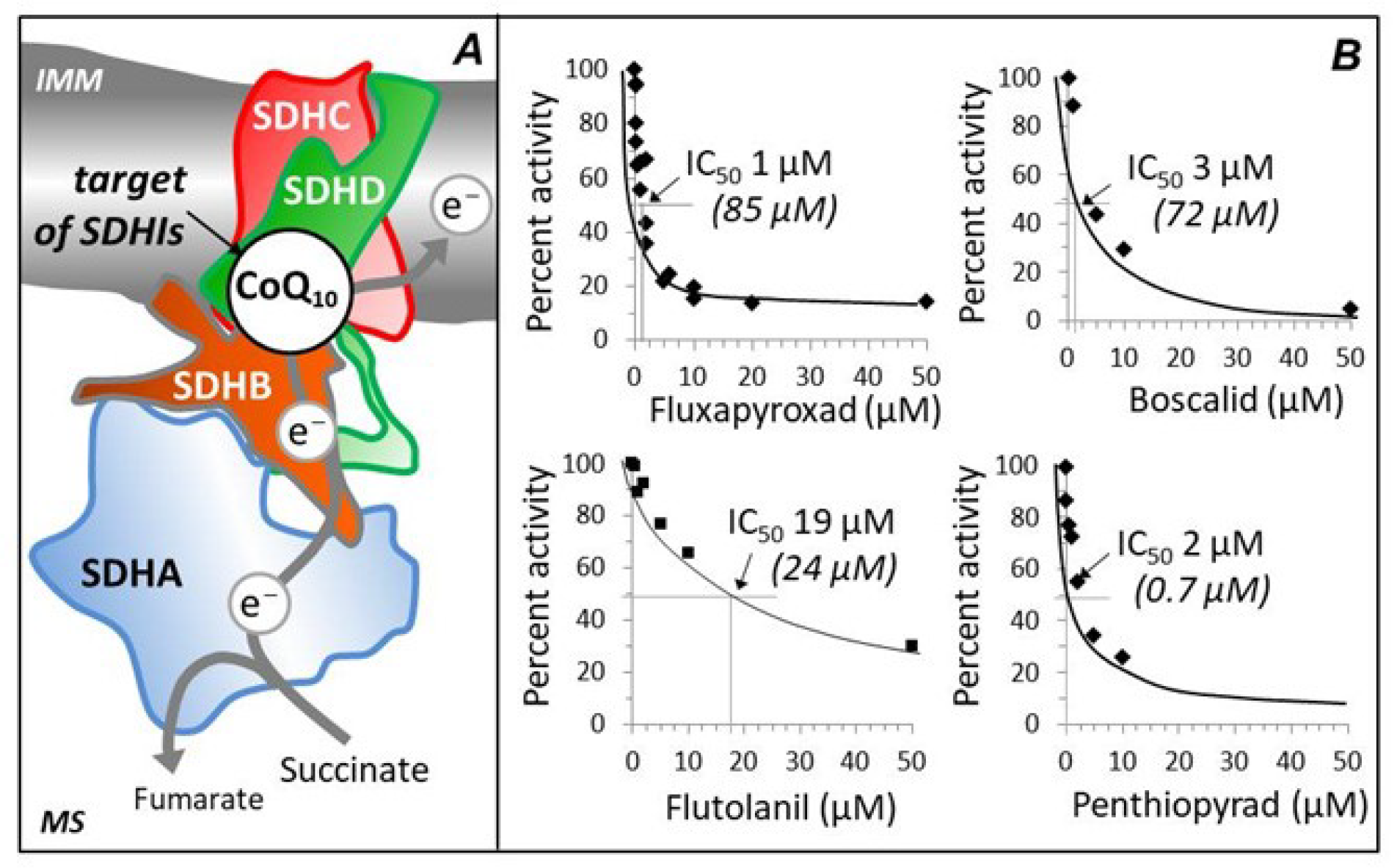
The succinate dehydrogenase and the SDHIs. **A**, a schematized view of the succinate dehydrogenase (SDH) in the inner mitochondrial membrane showing the ubiquinone (CoQ) reduction site involving three (SDHB-D) of the four subunits (SDHA-D) of the enzyme. Outlined by a dotted line, the site is supposedly the target of the SDHIs; **B**, The IC_50_ of various SDHIs toward the malonate-sensitive succinate cytochrome *c* reductase (SCCR) activity from human embryonic kidney (HEK) 293 cells. This SDH-dependent activity was selected because it does not require the addition of any exogenous quinone susceptible to alter the sensitivity to SDHIs. For each SDHI studied, the value of the IC_50_ measured with the enzyme from the earthworm, *Lumbricus terrestris*, is indicated in italics between brackets. Under all conditions (absence or presence of inhibitor), SCCR activity was measured as described (*1*) using 0.97 µg/ml and 2.36 µg/ml mg protein for the HEK293 cells (frozen cells, twice) and earthworm (head homogenate, low speed supernatant, 1 500 *g*) respectively, resulting (in the absence of any inhibitor, *i.e.* 100% activity) in a SCCR activity (n=4) of 75.9 ± 6 and 5.2 ± 0.7 nmol/min/mg protein, for the HEK293 cells and earthworm respectively. IMM, Inner Mitochondrial Membrane; MS, Matrix Space.

Altogether these observations were an incentive to study the effect of some of the available SDHIs on the human SDH (Table I). To our surprise, these studies revealed that the human enzyme is readily inhibited by these molecules with IC_50_ in the micro molar range. In parallel, we carried out a similar study on earthworm SDH revealing, depending on the considered molecule, varying degrees of sensitivity to the studied SDHIs as compared to human, suggestive of predictable quantitative difference according to species. Having for some of us dedicated our lives to fight the severe conditions resulting in human from a defect in SDH, it seems quite puzzling that inhibitors precisely targeting this enzyme could be so widespread in nature. The danger represented by recurrent presence in the food of these inhibitors, even at low dose, might even be higher for persons with decreased or abnormal activity of the succinate dehydrogenase, as established in the patients suffering some genetic diseases (i.e. Friedreich ataxia (*7*), Barth syndrome (*8*), Huntington’s disease (*9*)) or presenting with asthenozoospermia (*10*).

As recently reported (*11*), a high proportion of the pesticides in commerce have never been adequately tested for safety or toxicity. In that way, our concern is reinforced by the assumption that the cytotoxicity and genotoxicity tests that have possibly been used to assess toxicity of these compounds in human materials might have been inappropriate to properly evaluate their potential dangerousness. As previously noticed, cells with impaired SDH function can grow normally and standard genotoxicity tests are not designed to identify modifications of cellular epigenetics such as a DNA hypermethylated phenotype which may only appear after 15 cell passages in culture (*12*). Of note, despite repeated attempts, a rodent model for SDH-associated tumorigenesis is still lacking suggesting that these animals can hardly be used as models for teratogenicity resulting from succinate accumulation (*13*). Thus, similarly to endocrine disruptors and low-dose and mixture exposure to chemicals (*14*), the risk assessment framework for SDHIs should be urgently revisited before further distribution of existing products or introduction of new molecules on the market.

Now, especially after the impressive decline of insect populations in Western Europe reported last year (15), the media have taken hold of the subject, popularizing the recent declarations of renowned scientists, such as Hubert Reeves, expressing their major concerns about the consequence of earthworm soils population depletion, or the work carried out by two networks monitoring birds in the French countryside supported by the *Muséum national d’Histoire naturelle* (MNHN) and the *Centre National de la Recherche Scientifique* (CNRS) reporting the virtual disappearance, especially from 2008-2009, of several wild field birds (information at the front page of *Le Monde* and *The Guardian*). In all cases, the large-scale use of pesticides has been incriminated as being the major cause of these dramatic disruptions of our environment.

For some of them, pesticides have one or more undefined targets, an ambiguity behind which the supporters of their use attempt to shelter. Practically, they drive behind a fraction of the public opinion and, this latter in its path, a part of political decision-makers. Such is not the case for the SDHIs whose target and specificity are not disputed by manufacturers, users or scientists as we are. As a result, the danger we highlight regarding the inhibition of SDH on the basis of unambiguous scientific works should be taken seriously. These works cannot be suspected to be partisan since they have been mostly acquired before the marketing of the SDHIs and published in prestigious scientific journals. Difficulty to ban pesticides is all the more true as the economic stakes of changing agriculture practice is potentially considerable, requiring resolution and long term determination from our political decision-makers. However, the potent deleterious effects of the SDHIs on SDH from different species, including human, let us to think that the precautionary principle should lead to reconsider the widespread release of SDHIs in the environment. Based on the danger represented by SDHIs they should urgently be added to the list of pesticides which utilization should be either proscribed or restricted to strict necessity. Simultaneously the several years and tedious studies which are required to quantitatively estimate the risk they represent for human, and the fauna exposed to these drugs should now be launched. While awaiting the long and tedious studies required to estimate quantitatively the risk they represent for human, and the fauna exposed to these drugs, their use should be restricted to minimum

## References

1. T. Bourgeron et al., Nat Genet 11, 144 (Oct, 1995).

2. B. E. Baysal et al., Science 287, 848 (Feb 4, 2000).

3. A. J. Gill et al., Pathology 45, 689 (Dec, 2013).

4. J. J. Briere et al., Hum Mol Genet 14, 3263 (Nov 1, 2005).

5. E. Letouze et al., Cancer Cell 23, 739 (Jun 10, 2013).

6. C. Pouchieu et al., Int J Epidemiol, (Nov 9, 2017).

7. A. Rotig et al., Nat Genet 17, 215 (Oct, 1997).

8. J. Dudek et al., EMBO Mol Med 8, 139 (Feb 1, 2016).

9. S. J. Tabrizi et al., Ann Neurol 45, 25 (Jan, 1999).

10. R. Tomar, A. K. Mishra, N. K. Mohanty, A. K. Jain, Am J Reprod Immunol 68, 486 (Dec, 2012).

11. P. J. Landrigan et al., Lancet, (Oct 19, 2017).

12. S. Turcan et al., Nature 483, 479 (Feb 15, 2012).

13. C. Lepoutre-Lussey et al., Mol Cell Endocrinol 421, 40 (Feb 5, 2016).

14. W. H. Goodson, 3rd et al., Carcinogenesis 36 Suppl 1, S254 (Jun, 2015).

15. C. A. Hallmann et al., PLoS One 12, e0185809 (2017).

